# NDC1 is necessary for the stable assembly of the nuclear pore scaffold to establish nuclear transport in early C. elegans embryos

**DOI:** 10.1101/2021.07.17.452264

**Authors:** Michael Sean Mauro, Gunta Celma, Vitaly Zimyanin, Kimberley H. Gibson, Stefanie Redemann, Shirin Bahmanyar

## Abstract

Nuclear pore complexes (NPCs) are large protein assemblies that facilitate transport of macromolecules across the nuclear envelope (NE) [1, 2]. How thousands of NPCs rapidly assemble after open mitosis to form a functional NE is not known. Recruitment of the Nup107-160 outer ring scaffold to chromatin initiates NPC assembly. The Nup53/93 complex bridges the outer ring to the central channel to form a functional pore [3–6]. Nup53 interacts with the conserved transmembrane nucleoporin Ndc1; however, how Ndc1 contributes to post-mitotic NPC assembly is unclear [7–9]. Here, we use *C. elegans* embryos to show that the timely formation of a functional NE after mitosis depends on Ndc1. Endogenously tagged Ndc1 is recruited early to the reforming NE and is highly mobile in the nuclear rim. 3D analysis of post-mitotic NE formation revealed a decreased NPC density in NEs of *ndc1* deleted embryos – continuous nuclear membranes contained few holes where assembling NPCs are normally located. Nup160 is highly mobile in NEs depleted of Ndc1 and outer ring scaffold components are less enriched at the rim. When both *ndc1* and *nup53* are absent, nuclear assembly fails. Together, these data show that Ndc1 dynamically associates with the NE and promotes stable association of the outer ring scaffold in the NE to facilitate NPC assembly after open mitosis. Furthermore, Ndc1 and Nup53 function in parallel to drive nuclear assembly. We propose that Ndc1 is a dynamic membrane adaptor that helps recruit and promote the self-assembly of the nuclear pore scaffold to drive post-mitotic NPC assembly.

## Results and Discussion

A previous genome-wide high content RNAi-screen using differential interference contrast (DIC) to film early *C. elegans* embryos showed that RNAi-depletion of *ndc1* results in pronuclei that are smaller in size than in control embryos [10]. We used CRISPR-Cas9 gene editing to delete the *ndc1* gene locus and eliminate *ndc1* mRNA expression (Figure S1A and S1B; *ndc1Δ*). *ndc1Δ* embryos contained small pronuclei (Figure 1B) that were similar in size to those RNAi-depleted for *ndc1* (Figure S1C). Embryos produced from homozygous *ndc1Δ* worms had a range of lethality (40% - 100%, average ± S.D. = 61% ± 21, Figure 1C), that was similar but more severe than embryos produced from a mutant strain of *ndc1 (ndc-1^tm2886^)* in which 50% of the *ndc1* gene coding region is deleted (range 10% - 80%, average ± S.D. = 48% ± 25%, Figure S1D and S1E; [11]). The size of *ndc1Δ* embryos, but not *ndc1* RNAi-depleted embryos, appeared smaller suggesting a potential defect in germline development that may explain the embryonic lethality that was not observed by RNAi-depletion of *ndc1* (Figure 1B, S1C, and S1F). The fact that both *ndc1Δ* and RNAi-depletion of *ndc1* consistently result in small pronuclei suggested that Ndc1 is necessary for a process that controls nuclear size in early embryos.

**Figure 1.**
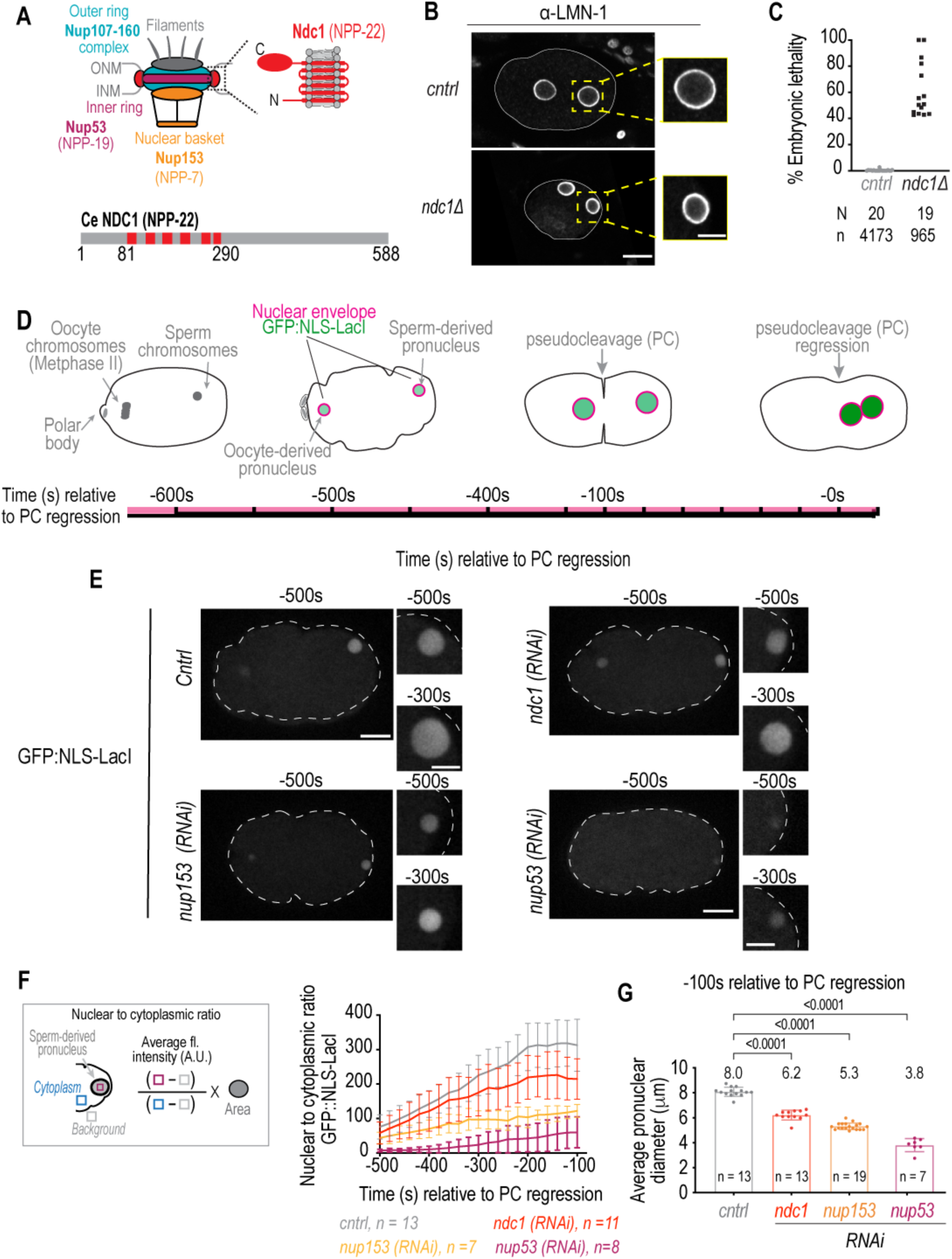
Smaller pronuclear size resulting from loss of Ndc1 corresponds to reduced nuclear import in early *C. elegans* embryos. **(A)** Schematic of an NPC (left). Schematic and domain organization of *Ce* Ndc1 (right, bottom) **(B)** Fixed overview and magnified images of *C. elegans* embryos immunostained for lamin for indicated conditions. Scale bars, 10 μm and 5 μm for magnified images. **(C)** Plot of percentage embryonic lethality for indicated conditions. N = # of worms. n = number of embryos. **(D)** Schematic of stereotypical nuclear events relative to nuclear import in early embryos in the *C. elegans* zygote, with pseudocleavage (PC) regression used as a reference time point. Time is in seconds. **(E)** Confocal overview and magnified images of embryo from a time lapse series of GFP:NLS-LacI in indicated conditions. Scale bars, 10 μm for overview image and 5 μm for magnified images. **(F)** Schematic for equation to calculate nuclear to cytoplasmic ratio of GFP::NLS-LacI (left) and corresponding plot for indicated conditions (right). Average ± S.D. is shown. **(G)** Pronuclear diameter for indicated conditions at indicated timepoint. Average ± S.D. is shown. n = # of embryos. A two-way ANOVA was used to determine statistical significance between control and each RNAi condition. p-values all < 0.0001.

The rate of nuclear import predicts the rate of nuclear expansion and nuclear size [12–15]. The rapid and stereotypical early divisions of *C. elegans* embryos provides an opportunity to quantitatively monitor nuclear import from the onset of nuclear formation until nuclear permeabilization in mitosis (Figure 1D; [16]). Fertilization of the oocyte by haploid sperm initiates two rounds of meiotic chromosome segregation. The sperm chromatin is devoid of a nuclear envelope (NE) at the time of fertilization. After anaphase of meiosis II, components in the oocyte cytoplasm are recruited to the sperm-derived pronucleus to form a functional NE (Figure 1D). Transport competent pronuclei then expand ∼30-fold in volume prior to the onset of nuclear permeabilization, which follows pronuclear meeting and regression of the pseudocleavage (Figure 1D; [17]). Nuclear import monitored by a GFP reporter fused to a nuclear localization (GFP:NLS-LacI, hereafter referred to as GFP:NLS) shows nuclear accumulation of GFP fluorescence throughout the time course of nuclear expansion (Figure 1D-1F). RNAi-depletion of Ndc1 results in lower levels of nuclear accumulation of the GFP:NLS reporter (Figure 1E-1F, Video S1). Nuclear import is more severely impaired in embryos depleted of the nuclear basket component Nup153 (*npp-7*) and the essential inner ring component Nup53 (*npp-19*) (Figure 1E-1F, Video S1; [18–22]). The average diameter of pronuclei is ∼8 μm ∼-100 s relative to pseudocleavage regression in control embryos (Figure 1G) and the diameter of pronuclei at the same timepoint in each RNAi condition scales with the degree of severity in defective nuclear import for that condition (6.2 μm for *ndc1(RNAi)* versus 5.3 μm for *nup153(RNAi)* and 3.8 μm for *nup53(RNAi)*; Figure1F and 1G). Thus, reduced nuclear import likely causes smaller pronuclei to form in the absence of Ndc1 suggesting that Ndc1 serves a specific function to promote nuclear transport in early *C. elegans* embryos.

To determine if Ndc1 promotes nuclear transport by contributing to the assembly of the NE, we first analyzed the dynamics and localization of Ndc1 during the first mitotic division. Ndc1 fused to mNeonGreen at its C-terminus using CRISPR-Cas9 gene editing (Ndc1^en^:mNG, Figure S2A and S2B) is enriched at the NE and appears functional given the normal pronuclear size in these embryos (Figure S2B, S2C, and Video S2). Ndc1^en^:mNG also localizes to the ER as well as punctate structures throughout the cytoplasm that disperse into the ER upon entry into mitosis (Figure S2C). Upon mitotic exit, Ndc1^en^:mNG accumulates at the reforming NE with similar kinetics as the inner NE protein LEM-2 (LEM-2:mCh; Figure 2A, 2B, and Video S3) and coincident with the appearance of signal from the fluorescent ER maker SP12:GFP around segregated chromosomes (Figure S2D). The early recruitment of Ndc1 to the post-mitotic NE suggested a potential role for Ndc1 in establishment of nuclear transport.

**Figure 2.**
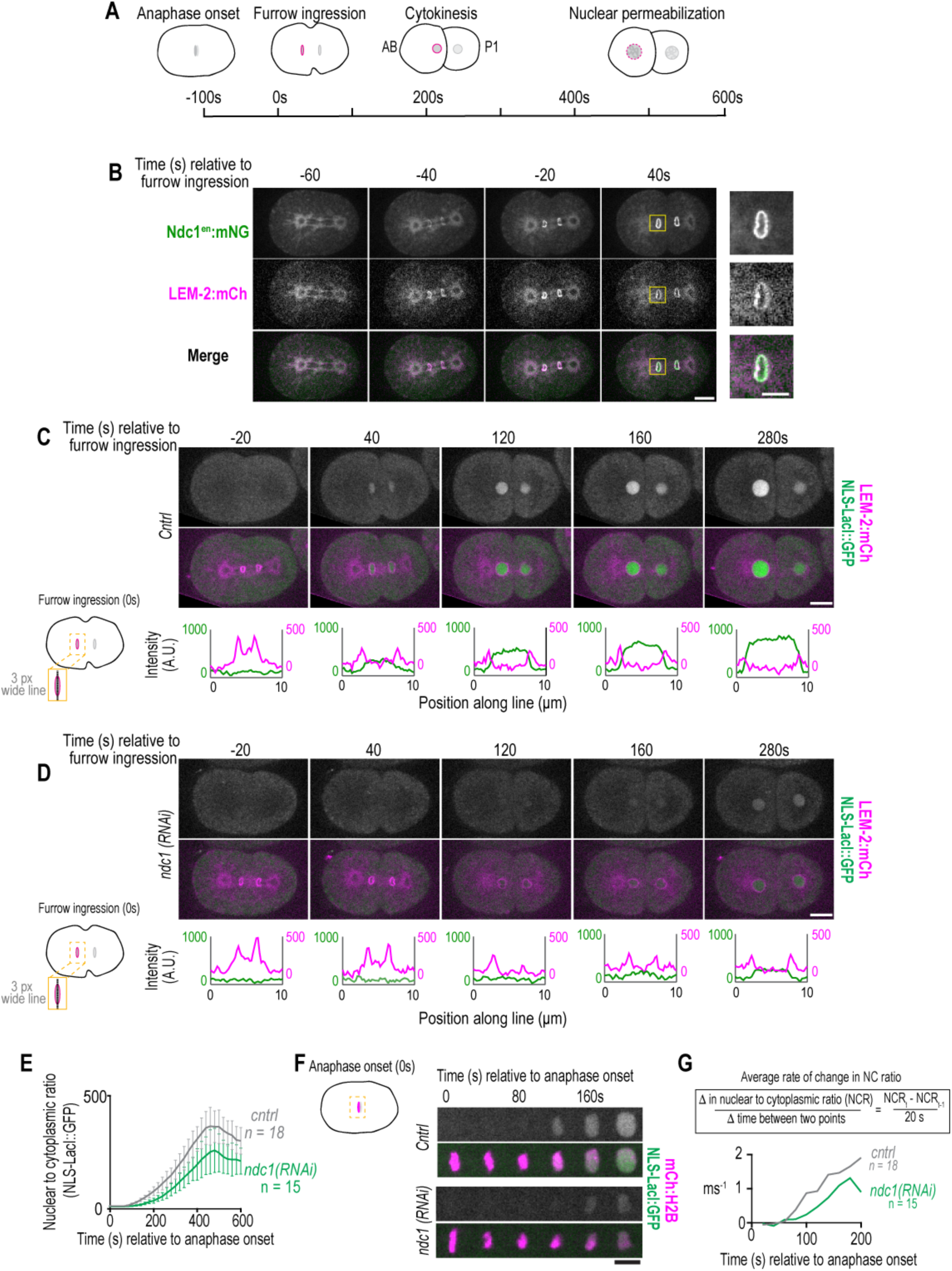
Ndc1 is necessary for timely formation of a transport competent nucleus after mitosis. **(A)** Schematic of the first mitotic division in *C. elegans* embryos elative to anaphase onset and initiation of furrow ingression. All measurements of 1 to 2 cell stage embryos are done on the AB nucleus (NE is highlighted magenta). **(B)** Confocal images from time series of mitotic nuclear formation relative to furrow ingression with indicated markers. **(C-D)** (Above) Confocal images from time series mitotic nuclear formation with indicated markers for indicated conditions. (Below) Line scans measuring background-corrected fluorescent intensities as indicated in schematic for each time point and fluorescent marker. Scale bars, 10 μm and 5 μm for magnified images. **(E)** Plot of nuclear to cytoplasmic ratio of GFP:NLS-LacI for indicated conditions. n = # of embryos. Average ± S.D. is shown. **(F)** Confocal images of chromosome region from time series relative to anaphase onset with indicated markers and in indicated conditions. Scale bar, 5 μm**. (G)** Equation and plot of rate of change in the nuclear to cytoplasmic ratio of GFP:NLS-LacI between each time point in the first 200 s after anaphase for the indicated conditions. Average is shown.

We next analyzed the kinetics of NE formation relative to the establishment of nuclear transport directly after anaphase onset. The NE rapidly forms around daughter nuclei ∼100 s after anaphase onset, coincident with the ingression of the cytokinetic furrow and followed by expansion of daughter nuclei (Figure 2A and 2C). In control embryos, the GFP:NLS fluorescence signal appears in the nucleus ∼40 s following initiation of furrow ingression, which is ∼60 – 80 s following the initial presence of a complete nuclear rim marked by LEM2:mCh (-20 s, Figure 2C). GFP:NLS continues to accumulate in the nucleus as nuclei expand (Figure 2C). In the absence of *ndc1*, LEM-2:mCh forms a nuclear rim with normal timing, however the GFP:NLS fluorescence signal is only detectable in the daughter pronuclei at ∼120 s following initiation of furrow ingression, which is ∼80 – 100 s later than in control embryos (Figure 2D). In addition to the later appearance of nuclear GFP, the nuclear to cytoplasmic ratio of the GFP:NLS fluorescence signal (see Figure 1F) was lower in *ndc1* RNAi-depleted embryos throughout the rest of the cell cycle (Figure 2E, Video S4).

To examine if the decreased levels of nuclear GFP:NLS fluorescence signal after anaphase onset in *ndc1* RNAi-depleted embryos (Figure 2E and 2F) result from a delay in establishment of nuclear transport, we normalized each individual trace of the nuclear to cytoplasmic ratio to its maximum value (Figure S2E). These data revealed a shift in the onset of nuclear accumulation of GFP:NLS in *ndc1* RNAi-depleted embryos (Figure S2E). Plotting the difference between each time point for the average normalized fluorescence intensities further revealed a slower rate of change in the nuclear accumulation of GFP:NLS after anaphase onset in *ndc1* RNAi-depleted embryos (Figure 2G). These data indicate that loss of Ndc1 delays nuclear import upon NE reformation.

The early recruitment of Ndc1 to the nascent post-mitotic NE combined with the delay in nuclear import resulting from loss of Ndc1 indicates that Ndc1 is necessary for the timely establishment of nuclear transport upon NE formation.

To determine how Ndc1 establishes nuclear transport upon NE formation, we analyzed serial sections of embryos from electron tomograms of NE formation processed at the initiation of furrow ingression (Figure 3A and 3B; Videos S5, S6 and S7). We focused on nascent nuclear membranes wrapped around the outer edges of chromatin where NPCs first assemble, also known as the “non-core” region [23]. 3D analysis of the reforming NE revealed small and large gaps at this time point and those <100 nm were marked as potential NPCs (Figures 3A, 3B, S3A, and S3B; Videos S8 and S9; [5]). The NE in *ndc1Δ* embryos was mostly continuous and contained an average of 8.6 “NPC” holes per μm^2^ (n = 3 areas) (Figure 3B, S3B and Video S9). In contrast, the NE in a control embryo contained an average of 37 “NPC” holes per μm^2^ (n = 2 areas) and was more discontinuous (Figure 3A and 3C; Video S8). Thus, when *ndc1* is absent, nascent NEs of the “non-core” region are more continuous and on average contain ∼4.3 fold fewer holes that fit the dimensions of nascent NPCs [5]. These data together with the slower rate of nuclear import resulting from loss of Ndc1 suggested that Ndc1 promotes NPC assembly during NE formation. The lower density of NPCs may limit the rate of accumulation of nuclear cargo during early stages of NE formation since soluble cargo are also lost by diffusion across larger gaps that are not yet sealed [24].

**Figure 3.**
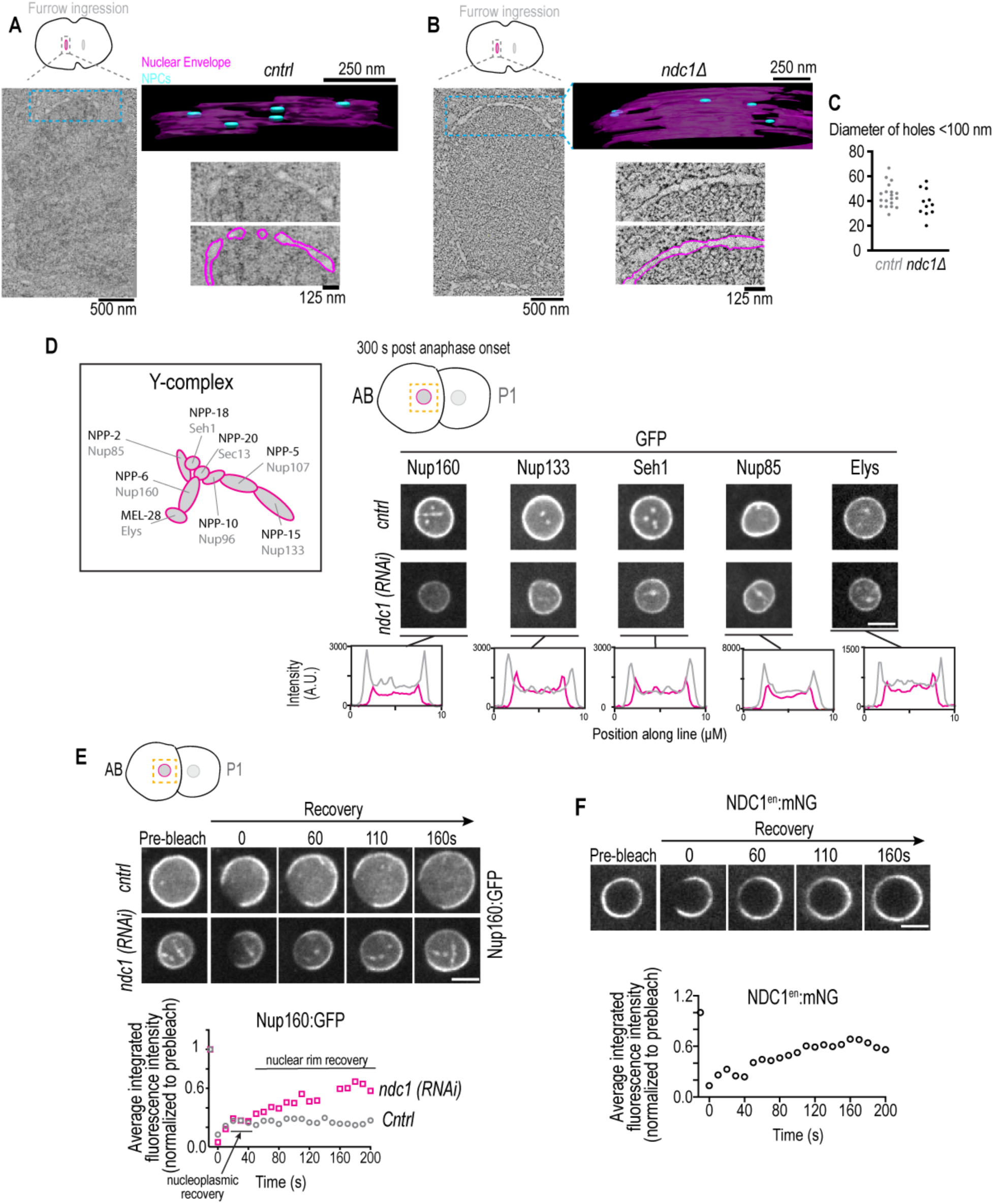
Ndc1 is mobile in the nuclear envelope and loss of Ndc1 results in fewer NPCs assembled on nascent NEs and a higher mobile pool of Nup160:GFP. **(A, B)** Overview images: z-slice from electron tomogram of nuclear formation timed relative to initiation of furrow ingression. 3D model: traced membranes (magenta) and NE holes < 100 nm (blue) for region shown as single z-slice in overview image. Magnified representative z slice traced and untraced from electron tomogram is shown. **(C)** Plot of diameters of NE holes < 100 nm. **(D)** Schematic of Y-complex with vertebrate (grey) and *C. elegans* (black) names shown. Magnified images of AB nucleus from confocal series for each indicated marker (above) and line scans measuring background-corrected fluorescent intensities (below) for each condition. **(E)** Confocal images from time lapse series of fluorescence recovery after photobleaching of Nup160:GFP for indicated conditions is shown (above). Plot of average fluorescence intensity of bleached region over time normalized to the prebleach intensity for each condition is shown (below). **(F)** Confocal images from time lapse series of fluorescence recovery after photobleaching of Ndc1^en^:GFP is shown (above). Plot of average fluorescence intensities of bleached region over time normalized to the prebleach intensity for each condition is shown (below). Scale bars, 5 μm.

The outer ring scaffold is essential for forming NEs that contain NPCs [6] and so we assessed whether loss of Ndc1 affects the localization of members of the outer scaffold complex (also known as the Y-complex [1, 2], Figure 3D). In early embryos, the Y-complex component Nup160:GFP localizes to kinetochores and to the nuclear rim (Figure S3C; [25]). Nup160:GFP also localizes to puncta throughout the cytoplasm that occasionally co-localize with Ndc1 puncta and disperse upon entry into mitosis (Figure S3D and S3E). *ndc1* RNAi-depleted embryos contained lower levels of Nup160:GFP at the nuclear rim (Figure 3D) and very few puncta formed in the cytoplasm (Figure S3E). Nup160:GFP puncta may represent preassembled nucleoporins known to be abundant in early embryos that contain excess nucleoporins [26]. The fact that they co-localize with Ndc1 and do not form in *ndc1* RNAi embryos, which also have lower levels of Nup160:GFP at the nuclear rim, suggested that Ndc1 may play a role in stabilizing scaffold components both in these cytoplasmic structures and in the NE.

The reduced nuclear rim signal of Nup160:GFP resulting from loss of *ndc1* likely reflects the lower density of NPCs in the NE. In addition to Nup160:GFP, line profiles of fluorescence intensities of GFP fusions to Nup133^NPP-15^, Seh1^NPP-18^, Nup85^NPP-2^, and Elys^MEL-28^ revealed that each component is approximately 50% less enriched at the nuclear rim in the absence of Ndc1 (Figure 3D; [25]). Immunostaining with antibodies that recognize Nup107^NPP-5^ and Elys^MEL-28^ confirmed lower endogenous levels of Y-complex nucleoporins at the nuclear rim in *ndc1Δ* early embryos (Figure S3F and S3G; [27, 28]). The total protein levels of Nup107 in *ndc1Δ* worms were similar to control worms (Figure S3H). Deletion of *ndc1* also results in lower levels of the endogenous inner ring component Nup53 at the nuclear rim (Figure S4A; [19]), but not its global protein levels (Figure S4B). Immunostaining of control and *ndc1* mutant worms with mAB414, a monoclonal antibody that recognizes FG-nucleoporins, did not show a consistent difference in fluorescence signal at the nuclear rim in early stage *ndc1* mutant embryos (Figure S4C), although prior work in late-stage *C. elegans* embryos showed that loss of *ndc1* decreases the levels of FG nucleoporins at the NE [11]. Total protein levels of mAB414 epitope-containing nucleoporins and the FG nucleoporins Nup96/98 was unchanged (Figure S4D and S4E;). Together, these results provide further evidence that Ndc1 is involved in assembly of the NPCs at the NE in early *C. elegans* embryos.

The fact that Ndc1 is an integral membrane protein localized to the pore membrane has led to the proposal that it serves as a membrane anchor for NPCs [8], although currently direct evidence for this does not exist. Fluorescence recovery after photobleaching (FRAP) revealed that ∼ 80% of the Nup160:GFP pool in the NE is immobile in control embryos (Figure 3E and Video S10; average mobile fraction ± SD = 0.20 ± 0.06, n = 8 nuclei), similar to what has been shown in mammalian cells [29]. In *ndc1 RNAi* depleted embryos there was a greater than 2-fold increase in the mobile fraction of Nup160:GFP in the NE indicating that Nup160:GFP is less stably incorporated without Ndc1 (Figure 3E; average mobile fraction ± SD = 0.47 +/- 0.12, n = 8 nuclei).

We reasoned that if Ndc1 serves as an anchor to the NPC to immobilize Nup160:GFP, then Ndc1 itself should be immobile at the NE. Instead, we found that Ndc1^en^:mNG is highly mobile in the NE (Figure 3F and Video S11; average mobile fraction ± SD = 0.59 ±- 0.09, t_1/2_ of 56.8 ± 26.3s, n = 8 embryos). This data is in line with recent data using metabolic labeling in budding yeast showing that Ndc1, unlike other transmembrane nucleoporins, is readily exchanged in the NPC [30]. Thus, Ndc1 may function as a dynamic membrane adaptor that promotes the stable association of outer ring scaffold components in the NE.

Ndc1 binds directly to Nup53, which links the inner and outer rings of the NPC to the central channel through its association with the Nup96/98 complex and Nup155 (Figure 4A and S4F; [7-9, 31, 32]). Prior work had suggested that Ndc1 recruits Nup53 to the assembling NPC [8] and that Ndc1 functions in NPC assembly through its interaction with Nup53 [7–9]. To determine if Ndc1 and Nup53 cooperate in post-mitotic NPC assembly in *C. elegans*, we utilized a strain carrying a partially functional mutant allele of Nup53 (*nup53^tm2886^)* that is missing a central region important for its dimerization (Figure S4G; [19]). The mutant Nup53(Δaa 217-286) protein is expressed at lower levels compared to wild type Nup53 (Figure S4B; [19]) and on average ∼ 50% of embryos produced from homozygous *nup53^tm2886^* worms can survive to hatching, as has been shown previously (Figure 4B; [19]). Live imaging of *nup53^tm2886^* one-cell embryos revealed that sperm pronuclei expand and establish import (Figure 4C), similar to *nup53* RNAi-depleted nuclei (Figure 1E-1G) but to a lesser extent than control and *ndc1* RNAi-depleted embryos (Figure 1E-1G). RNAi-depletion of *ndc1* strongly enhanced embryonic lethality in the *nup53^tm2886^* strain (Figure 4B) - these embryos completely failed to assemble a NE around sperm chromatin (Figure 4C and 4D). Sperm chromatin in *nup53^tm2886^* mutant embryos depleted of *ndc1* remains compacted and nuclear import is never established, even at -100 s relative to PC regression when GFP:NLS accumulates to detectable levels in the sperm-derived pronucleus of *nup53^tm2886^* embryos (Figure 4C and 4E).

**Figure 4.**
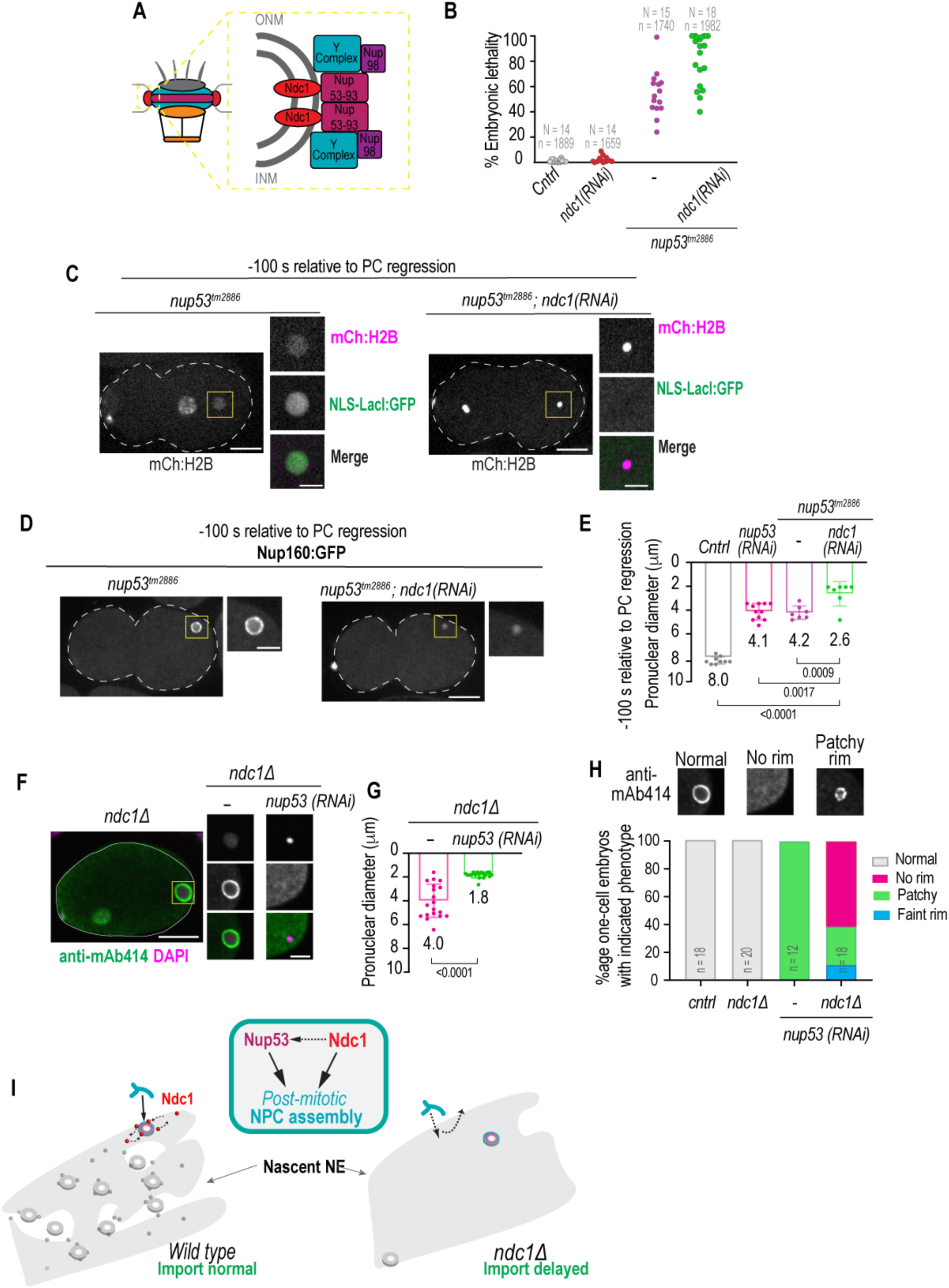
Ndc1 coordinates with Nup53 in nuclear assembly. **(A)** Schematic of NPC (left) and NPC subcomplex organization (right) is shown. **(B)** Plot of percentage embryonic lethality for indicated conditions. N = number of worms. n = number of embryos. **(C)** Confocal overview and magnified images of embryo from a time lapse series of indicated markers for indicated conditions. **(D)** Confocal overview and magnified images of embryo from a time lapse series of Nup160:GFP for indicated conditions. **(E)** Pronuclear diameter for indicated conditions at indicated time point. n = number of embryos. *cntrl* n = 9, *nup53(RNAi)* n = 11, *nup53^tm2886^* n = 7, and *nup53^tm2886^;ndc1(RNAi)* n = 7. **(F)** Fixed overview and magnified images of *C. elegans* embryos immunostained for mAb414 and DAPI in indicated conditions. Scale bars, 10 μm for overview image and 5 μm for magnified images. **(G)** Pronuclear diameter for indicated conditions at indicated time point. n = number of embryos. *ndc1Δ* n = 20 and *ndc1Δ;nup53(RNAi)* n = 18. **(H)** Magnified images of paternal pronucleus from fixed one-cell stage embryos immunostained with mAb414 for indicated conditions (top). Plot of mAb414 appearance surrounding chromatin under indicated conditions (bottom). A two-way ANOVA was used to determine statistical significance between indicated conditions. n = number of embryos. Scale bars, 10 μm for overview image and 5 μm for magnified images. **(I)** Schematic representation of NE formation with and without Ndc1. Schematic of a simplified genetic pathway highlighting the parallel contributions of Ndc1 and Nup53 to nuclear assembly.

In *nup53^tm2886^* mutant and *nup53* RNAi-depleted embryos, Nup160:GFP appears patchy at the sperm pronuclear rim indicating an abnormal distribution of NPCs (Figure 4D and S4H). In contrast, in *nup53^tm2886^* embryos RNAi-depleted of *ndc1*, the Nup160:GFP signal persisted on chromatin (Figure 4D) and the nuclear diameter at -100 s prior to PC regression was significantly smaller (Figure 4E; average diameter of 2.6 μm versus ∼4 μm in *nup53^tm2886^* or *ndc1(RNAi)* only conditions) indicating a failure in nuclear formation. These data show that Ndc1 is necessary for the recruitment of Nup160:GFP to the nuclear rim and assembly of nuclei, albeit in a defective manner, when Nup53 function is compromised.

Failure to assemble a NE was also observed in fixed *ndc1Δ* embryos RNAi-depleted for *nup53* (Figure 4F-4H). The mAB414 signal around chromatin was normal in *ndc1Δ* embryos and patchy in 100% of fixed one-cell stage embryos RNAi-depleted for *nup53* alone (Figure 4H and Figure S4I). The patchy Nup160:GFP (Figure 4D and S4H) and mAB414 (Figure 4H and S4I) signal resulting from loss of Nup53 suggests aberrant assembly of NPCs. Surprisingly, these NEs still support some nuclear import and expansion (Figure 4D, 4E and Figure 1E-1G). One-cell stage *ndc1Δ* embryos RNAi-depleted for *nup53* contained highly compacted sperm chromatin (Figure 4F and 4G) with little to no mAB414 signal surrounding the chromatin mass (Figure 4F and 4H). Together, these data provide evidence that Ndc1 and Nup53 function, at least in part, in parallel pathways to drive robust nuclear assembly in early *C. elegans* embryos.

Cumulatively, the results above demonstrate that Ndc1 is recruited early to the reforming NE after mitosis where it dynamically associates with the NE and serves to stabilize the outer ring scaffold to drive NPC assembly (Figure 4I). Our *in vivo* genetic analyses further demonstrate that the loss of *ndc1* strongly enhances the NE assembly defect resulting from loss of Nup53 (and vice versa) indicating that Ndc1 and Nup53 coordinate the assembly of a functional NE (Figure 4I).

We suggest that Ndc1 functions as a dynamic membrane adaptor that helps recruit and orient scaffold nucleoporins at membrane holes to promote their self-assembly and drive formation of NPCs. Electron tomography studies of the earliest stages of NE reformation revealed that assembly of nuclear pores initiates in 40 nm NE holes that then dilate upon scaffold assembly [5]. Interestingly, recent work has shown that preassembled Y-complex subunits associated with holes in the ER initiate NPC assembly after anaphase [33]. The fact that loss of *ndc1* results in nascent NEs that are continuous but with only a few holes <100 nm suggests that Ndc1 may be necessary for sculpting or maintaining these small holes that are for NPC assembly. Furthermore, the cytoplasmic puncta we observed with Ndc1 may be annulate lamellae, specialized membranes in the ER that are prominent in embryos and contain symmetric scaffold nuclear pores to store excess nucleoporins [26]. The fact that Nup160 cytoplasmic puncta sometimes colocalize with Ndc1 puncta in the ER and no longer form without Ndc1 suggests that Ndc1 may be involved in formation of these structures. However, we show that Ndc1 is highly dynamic in the membrane arguing against a role as a structural protein or immobile anchor at the pore membrane. More work needs to be done to understand the identity of these cytoplasmic puncta, their mechanistic relationship to Ndc1, and whether they are directly involved in NPC assembly in *C. elegans* embryos.

Given that there are still NPCs assembled in the absence of Ndc1, albeit at a significantly decreased density, indicates that scaffold nucleoporins can assemble without Ndc1, but perhaps at a slower rate. It is also possible that NPCs can also assemble through a parallel pathway to Ndc1. Supporting this idea, our work and work in other systems show that loss of Ndc1 and Nup53 inhibits NPC assembly [7–9, 21]. Nup53 has been shown to bind directly to membranes [21] and a region of Ndc1 that does not bind to Nup53 supports NPC assembly when Nup53’s membrane association region is deleted in *in vitro* assembled nuclei using *Xenopus* extracts [7]. This suggests that Ndc1 can drive NPC assembly independently of its interaction with Nup53 and supports our working model that these proteins function at least in part in parallel pathways to drive post-mitotic NPC assembly (Figure 4I). Future is required to determine how Ndc1 acts to immobilize the outer ring scaffold in the NE and how its role at the membrane coordinates with Nup53 to enable nuclear assembly.

## Acknowledgements

We thank Sarah Barger for helpful feedback on the manuscript and members of the Bahmanyar lab for helpful discussion. We thank Eric Hastie and Sarah Barger for help with cloning, the MBL Embryology course for insights into strain construction, Jackson Gordon for help with FRAP analysis. We thank Peter Askajar for sharing antibodies against various C elegans Nups, Arshad Desai, the Caenorhabditis Genetics Center and the Japanese knockout (KO) consortium for strains utilized in this study. This work was supported by an NSF CAREER Award to Shirin Bahmanyar (NSF CAREER 1846010), and NIH CMB Training Grant: T32GM007223-S1to Michael Mauro.

## Author Contributions

M.S. Mauro and S. Bahmanyar conceived the project. M.S. Mauro performed the majority of the experiments. G. Celma generated and filmed fluorescent strains used in this manuscript. V. Zimyanin and S. Redemann prepared samples by high-pressure freezing, generated electron tomograms, and joined the mutant tomogram. K.H. Gibson joined the wild type tomograms. M. S. Mauro generated 3D models of reconstructed tomograms. M.S. Mauro and S. Bahmanyar wrote the manuscript, with input from all authors. S. Bahmanyar supervised the project.

**The authors declare no competing interests**

---

**Figure S1.**
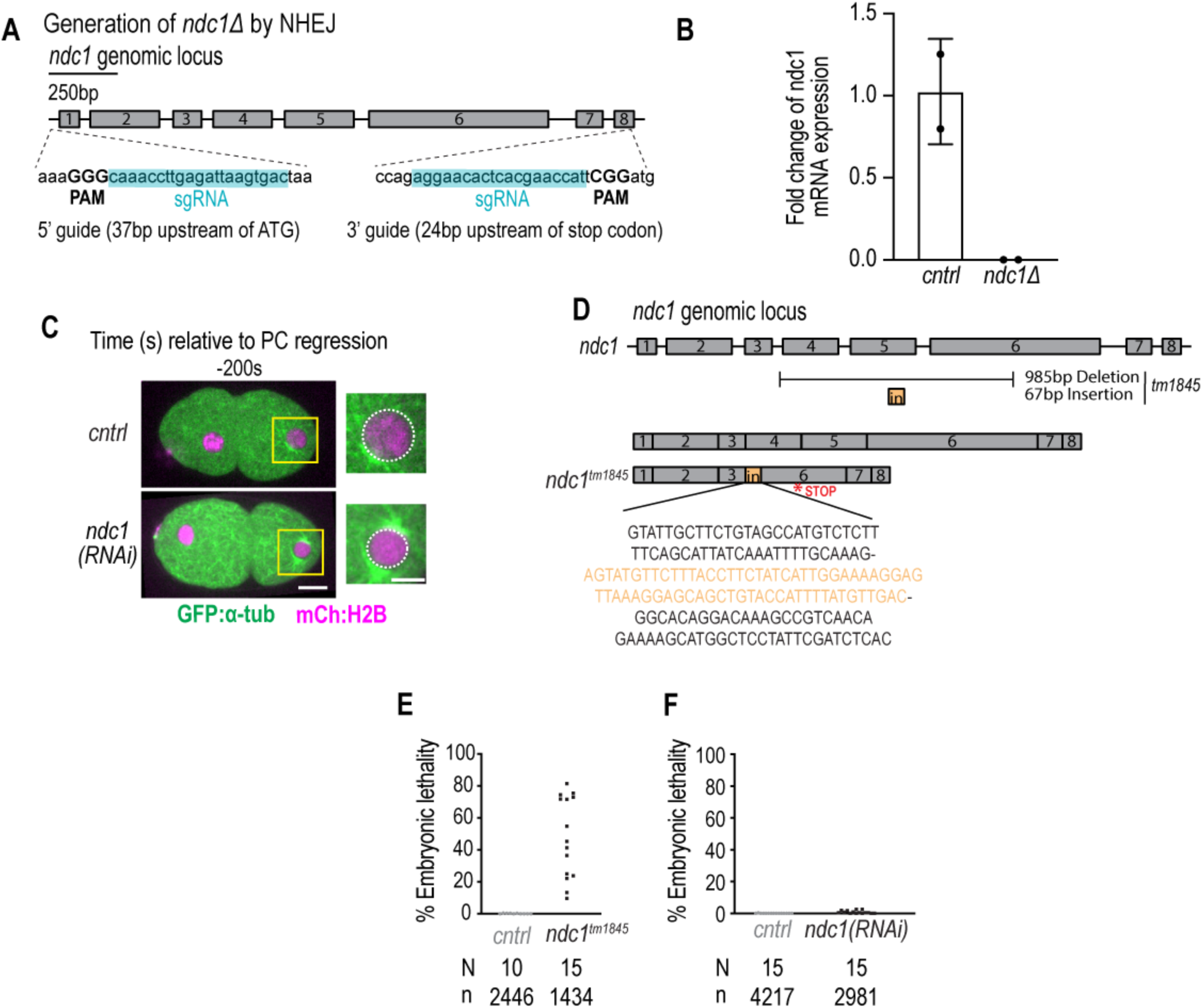
Generation and characterization of mutant *ndc1* alleles and RNAi-depletion of *ndc1*. *Related to Figure 1*. **(A)** Schematic of CRISPR guides to generate *ndc1* null allele. **(B)** Plot of *ndc1* mRNA fold change for control and *ndc1Δ* worms. Mean +/-SD. **(C)** Confocal overview and magnified images of one-cell stage embryo from a time lapse series of indicated markers for indicated conditions. Scale bars, 10 μm for overview image and 5 μm for magnified images. **(D)** Schematic of *ndc1* genomic locus (top) and *tm1845* allele (below). **(E-F)** Plots of percentage embryonic lethality for indicated conditions. N = number of worms. n = number of embryos.

**Figure S2.**
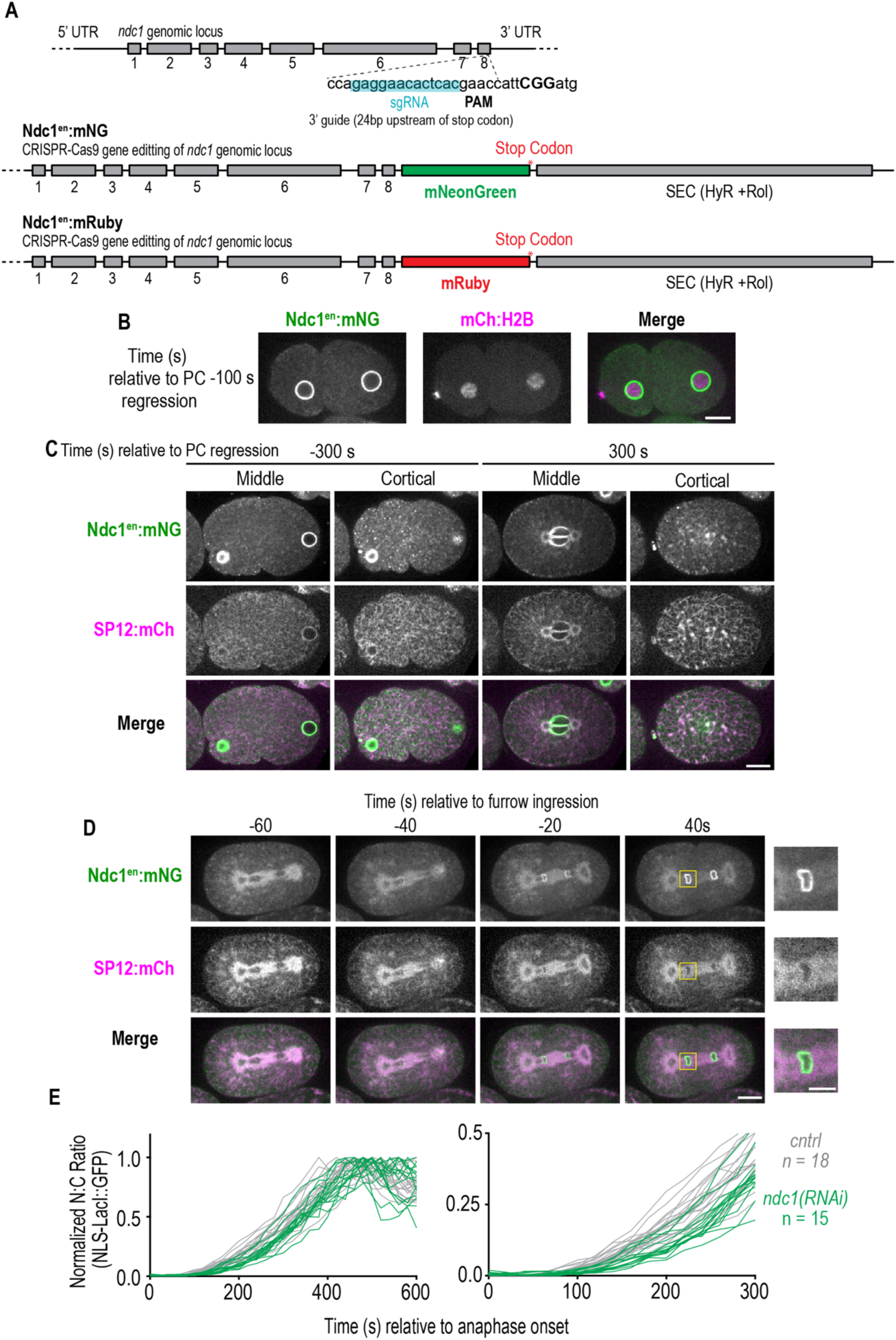
Ndc1^en^:mNG is recruited early to the nuclear rim and localizes to the ER and cytoplasmic puncta and nuclear transport is delayed in post-mitotic nuclei without Ndc1. *Related to Figure 2.* **(A)** Schematic of *ndc1* genomic locus and CRISPR guide to generate *ndc1^en^*:*mNG* and ndc1^en^:mRuby at its endogenous locus. **(B)** Confocal images from time series of one-cell stage embryo expressing Ndc1^en^:mNG and mCh:H2B. **(C)** Confocal images from time series of one-cell stage embryo expressing Ndc1^en^:mNG and SP12:GFP to mark the ER. (**D)** Confocal images from time series of two-cell stage embryo expressing Ndc1^en^:mNG and SP12:mCh during mitotic nuclear formation. Scale bars, 10 μm and 5 μm for magnified images. **(E)** Plot of normalized nuclear to cytoplasmic ratio of GFP:NLS-LacI for indicated conditions. n = number of embryos.

**Figure S3.**
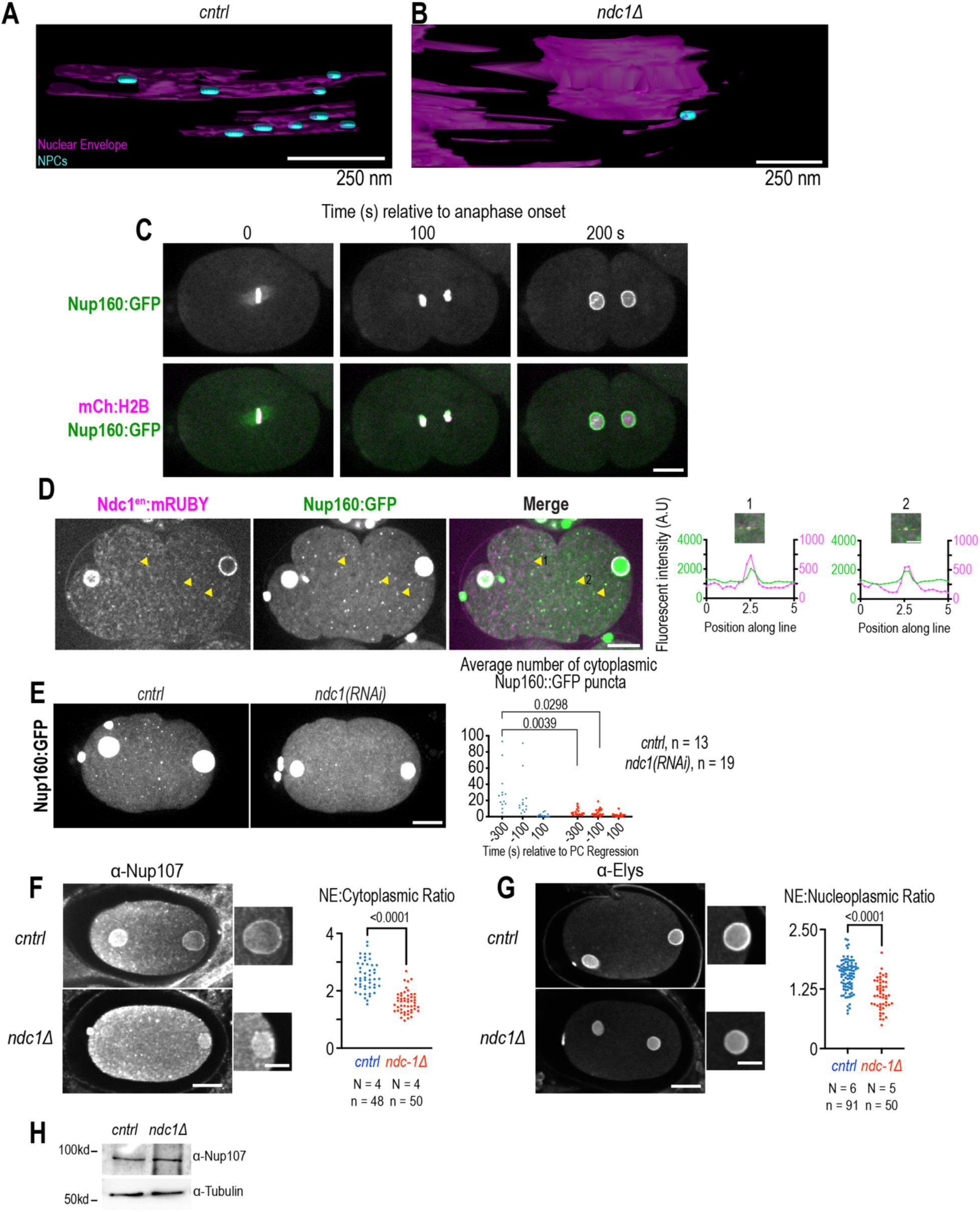
Lower nuclear pore density and levels of outer ring scaffold nucleoporins at the nuclear rim and cytoplasmic puncta resulting from loss of Ndc1. *Related to Figure 3.* (**A, B)** 3D model of tomograms of NE reformation showing membranes (magenta) and NE holes < 100 nm (blue) for indicated conditions. **(C)** Confocal images from time series of mitotic nuclear formation relative to anaphase onset with indicated markers in control embryos. **(D)** Confocal overview and magnified images of embryo from a time lapse series of Nup160:GFP and Ndc1^en^:mRuby (left). Yellow arrows indicate example puncta. Five μm line scans measuring background-corrected fluorescent intensities for each marker drawn across two representative cytoplasmic puncta with co-localized Ndc1 and Nup160 signal (right). Scale bars, 10 μm and 2.5 μm for magnified images. **(E)** Max projection confocal images of Nup160:GFP for indicated conditions (left). Plot of average number of cytoplasmic Nup160:GFP structures under indicated conditions at time points relative to PC regression (right). Scale bar, 10 μm. Students T-test was used to determine statistical significance. **(F)** Fixed overview and magnified images of *C. elegans* embryos immunostained with antibodies that recognize Nup107 for indicated conditions (left). Plot of NE to cytoplasmic ratio for normalized Nup107 fluorescence intensity (right). Scale bars, 10 μm and 5 μm for magnified images. **(G)** Fixed overview and magnified images of *C. elegans* embryos immunostained for antibodies against Elys^Mel-28^ for indicated conditions (right). Plot of NE to nucleoplasmic ratio for normalized Elys^Mel-28^ fluorescence intensity (right). Scale bars, 10 μm and 5 μm for magnified images. **(H)** Immunoblot of whole worm lysates probed for antibodies that recognize Nup107 and α-tubulin for indicated conditions.

**Figure S4.**
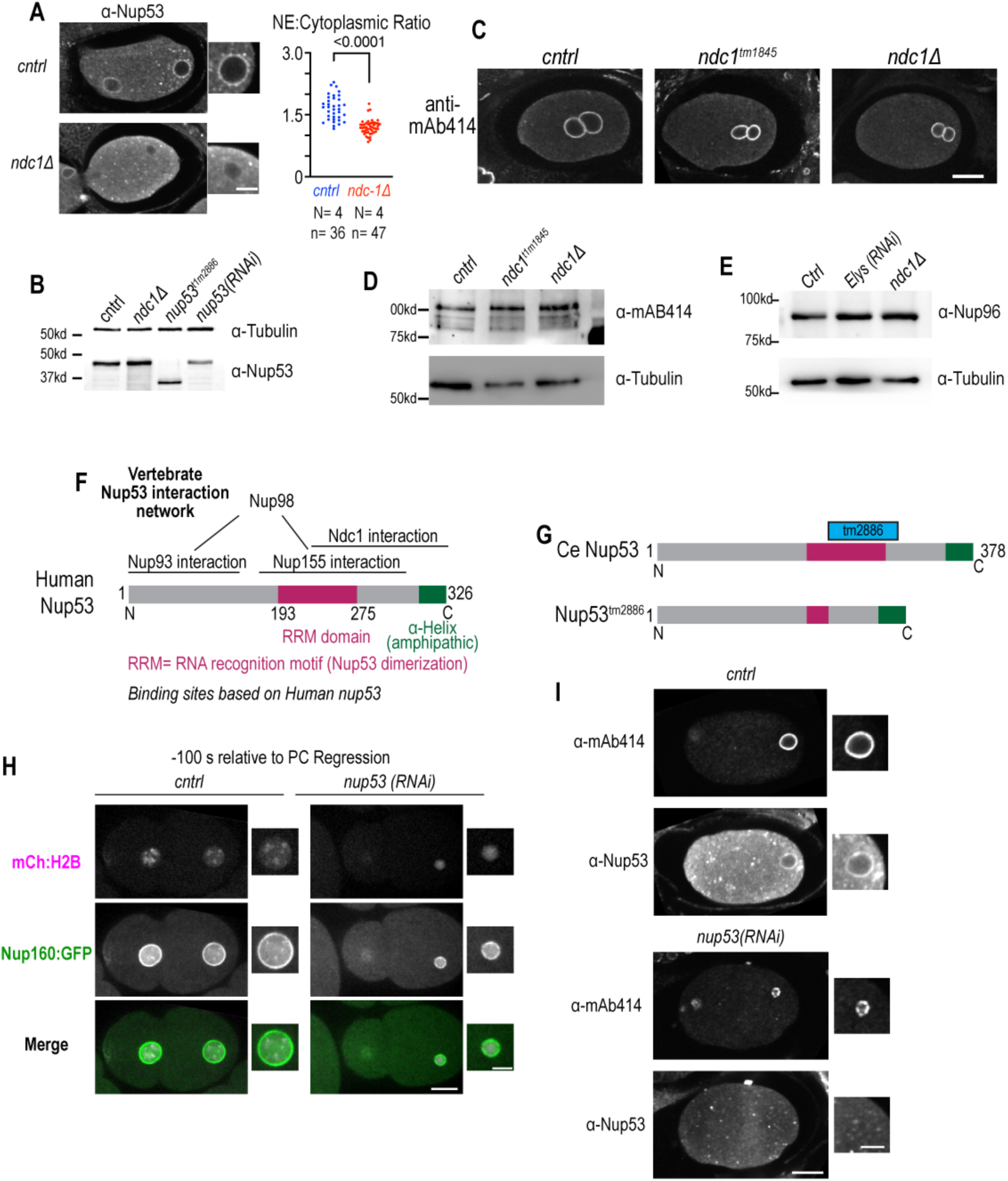
Characterization of levels and localization of nucleoporins in *ndc1* mutant and *nup53* mutant alleles. *Related to Figure 4.* **(A)** Fixed overview and magnified images of *C. elegans* embryos immunostained for antibodies that recognize Nup53 for indicated conditions (left). Plot of NE to cytoplasmic ratio for normalized Nup53 fluorescence intensity (right). **(B)** Immunoblot of whole worm lysates probed for antibodies that recognize Nup53 and α-tubulin for indicated conditions. **(C)** Fixed overview images of *C. elegans* oocytes immunostained for mAb414 for indicated conditions. Scale bars, 10 μm. **(D)** Immunoblot of whole worm lysates probed for mAb414 and α-tubulin for indicated conditions. **(E)** Immunoblot of whole worm lysates probed for Nup96 and α-tubulin for indicated conditions. **(F)** Schematic of Nup53 domain architecture and binding interactions based on human Nup53 binding sites. **(G)** Schematic of *C elegans* Nup53 domain architecture and the resulting mutant protein resulting from the *nup53^tm2886^* allele. **(H)** Confocal overview and magnified images of one-cell stage embryo from a time lapse series expressing Nup160:GFP and mCh:H2B for indicated conditions. Scale bars, 10 μm and 5 μm for magnified images. **(I)** Fixed overview and magnified images of *C. elegans* embryos immunostained for mAb414 and Nup53 for indicated conditions. Scale bars, 10 μm and 5 μm for magnified images.

## Key Resources

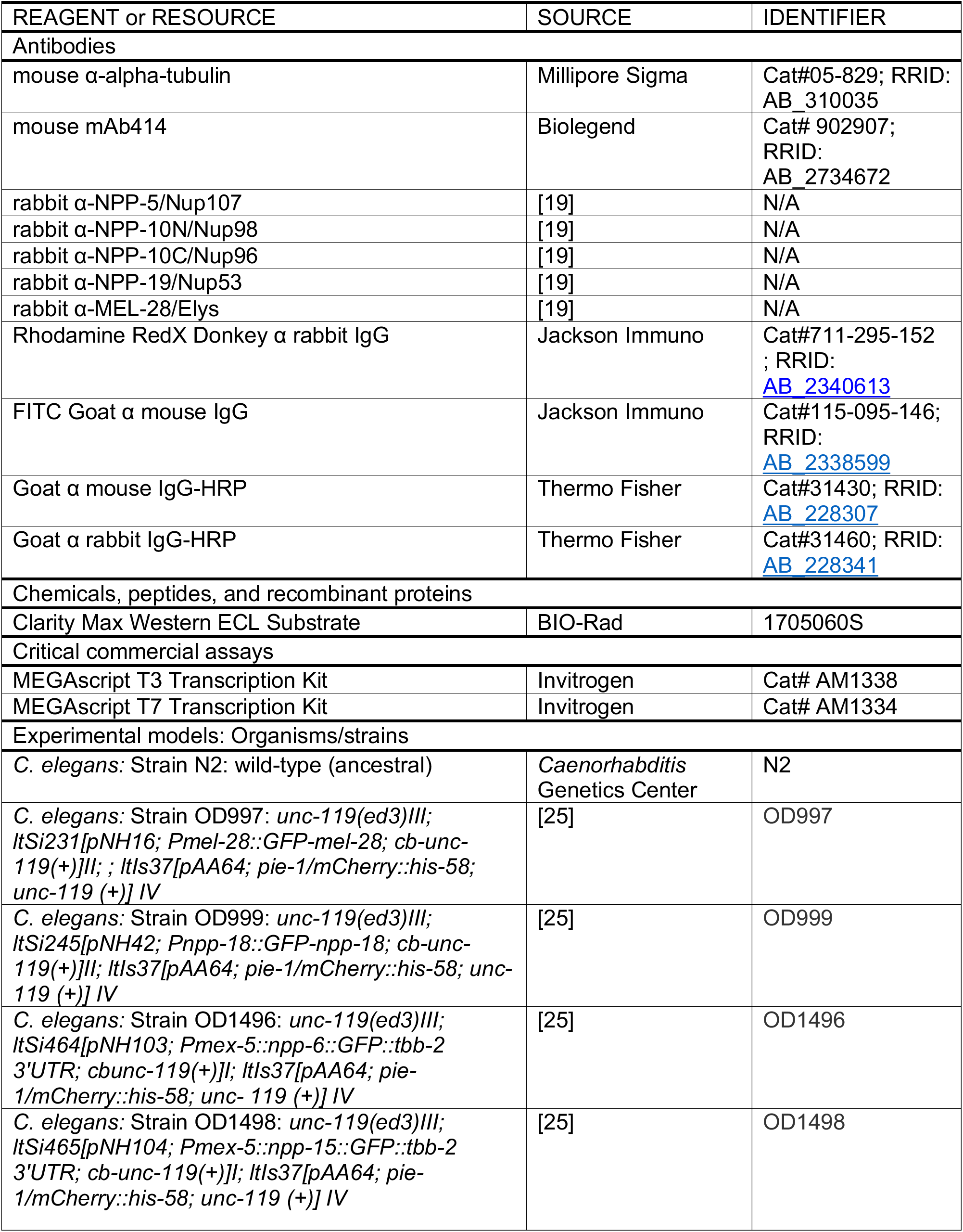
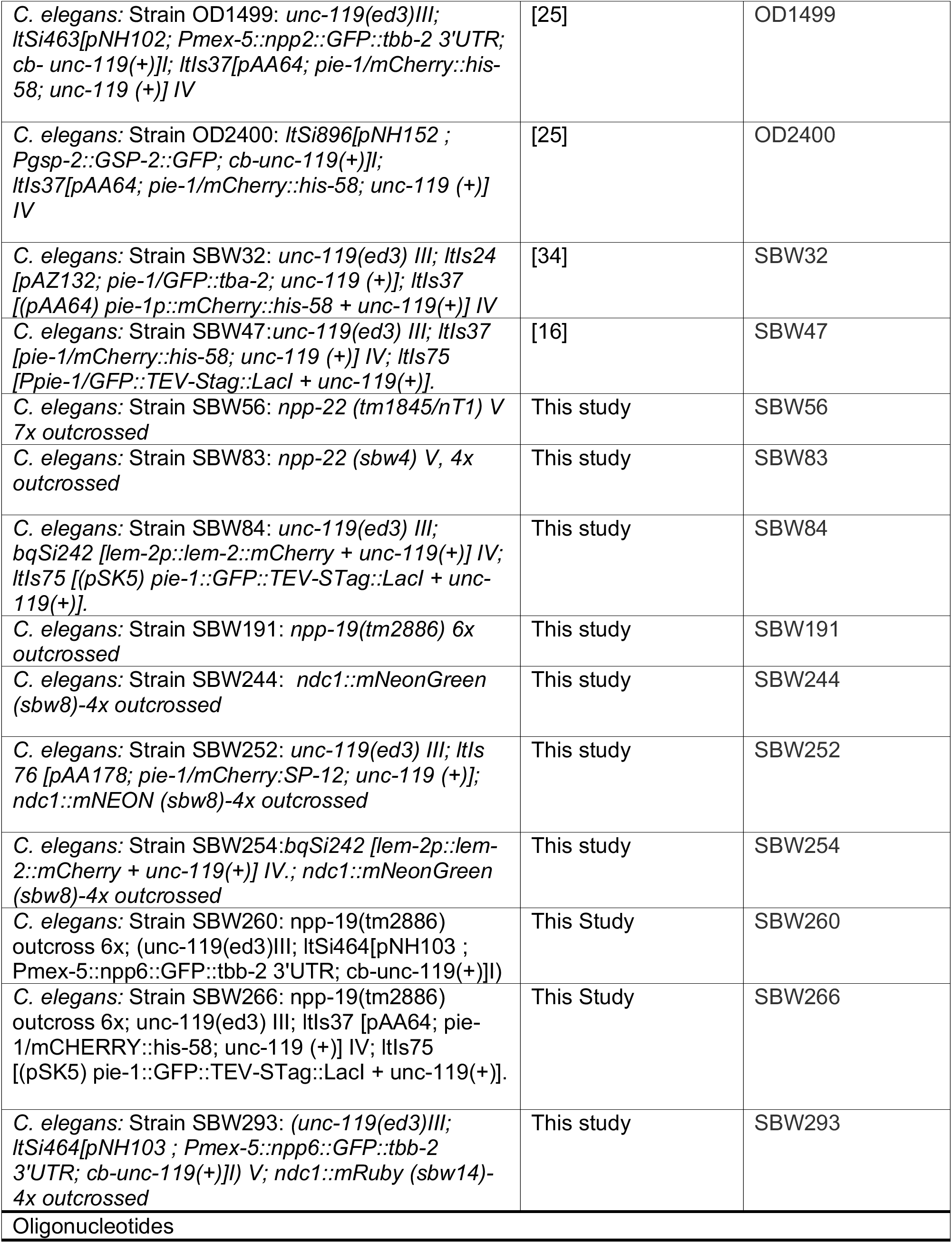
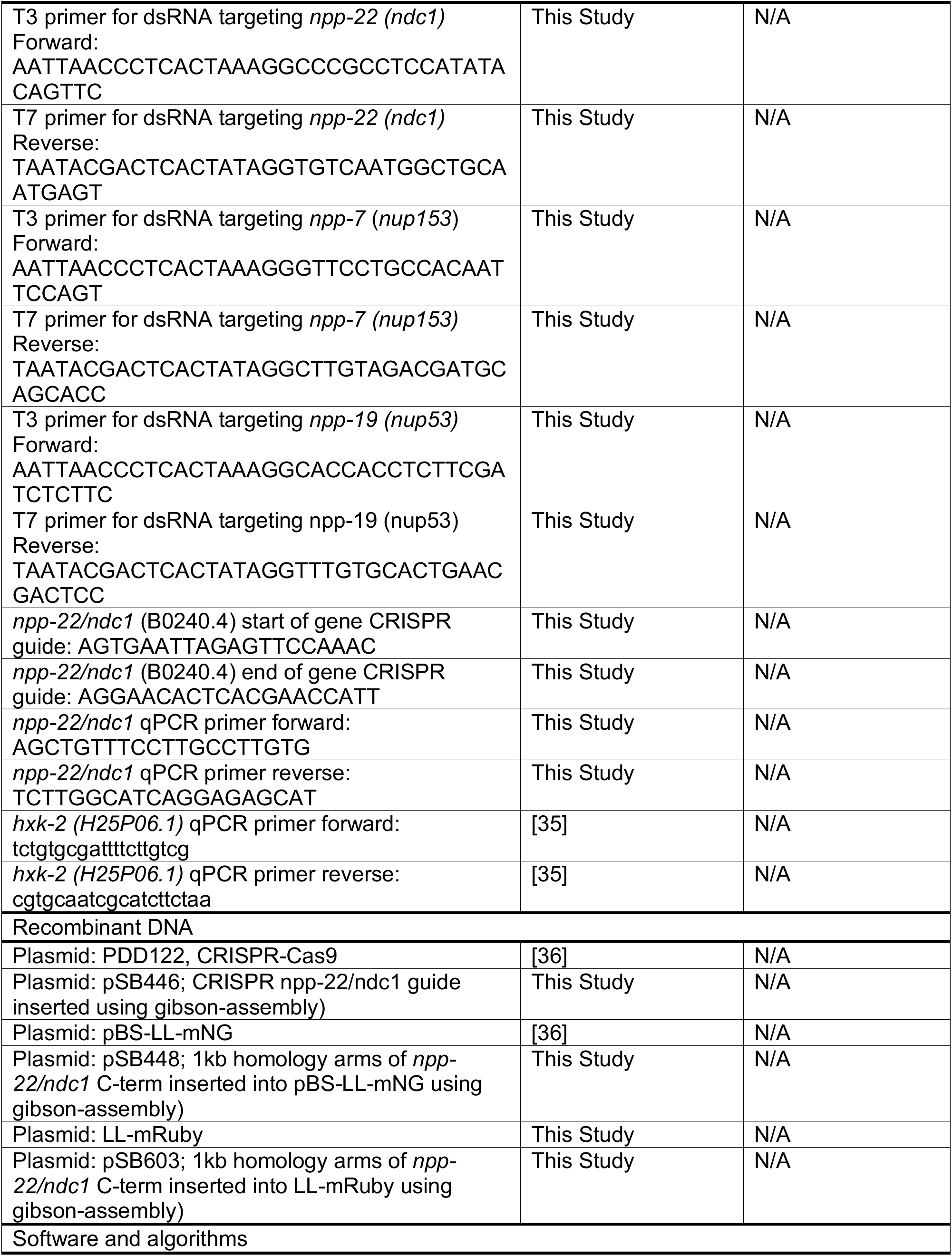
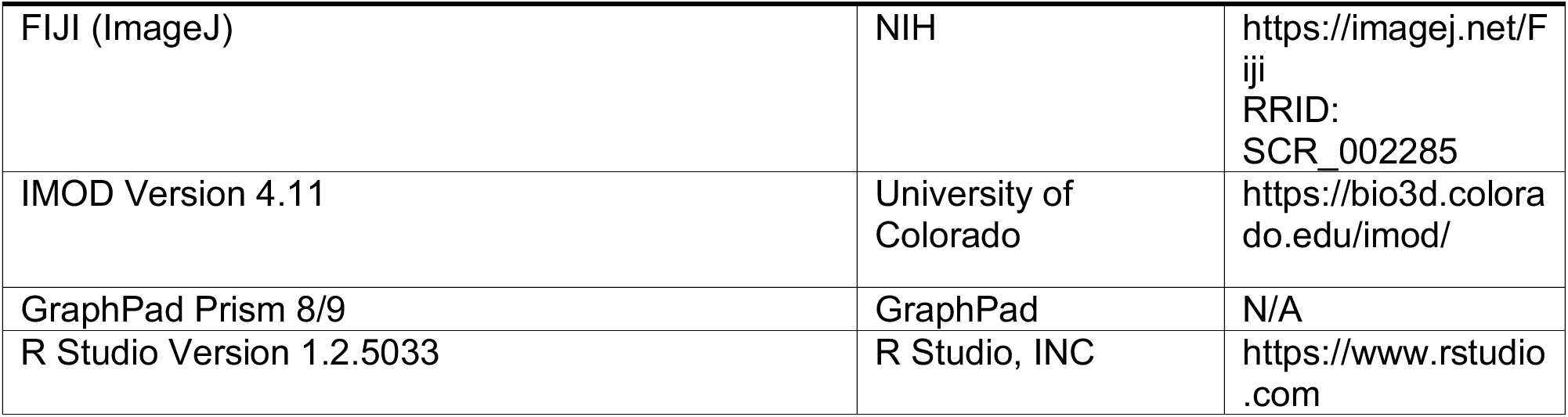

## RESOURCE AVAILABILITY

### Lead Contact

Further information and requests for reagents or resources should be direct to Shirin Bahmanyar.

### Materials availability

*C. elegans* strains and plasmids from this study are available from the Lead Contact upon request.

### Data and code availability

Raw data not included in this article are available from the corresponding author on request.

## EXPERIMENTAL MODEL AND SUBJECT DETAILS

### Strain maintenance and generation

The *C. elegans* strains used in this study are listed in the key resource table. Strains were maintained on nematode growth media (NGM) plates that were seeded with OP-50 *Escherichia coli.* Strains were grown at three possible temperatures: 15°C, 20°C and 25°C.

#### CRISPR-Cas9 deletion strain

The deletion for *ndc-1* (B0240.4) was generated using two CRISPR guides “crRNA”, which were obtained using IDT’s custom CRISPR guide algorithm (See **Fig. S1A** and key resource table for sequences). 1 μl of the purified crRNAs were annealed to 1 μl of trans-activating crRNA (tracrRNA) by incubating RNAs at 95 DC for two minutes in individual PCR tubes. A co-CRISPR crRNA was used for dpy-10. An injection mix with the following components and concentrations was set up and spun down for 30 minutes: at 4°C; *ndc-1* guide 1 (11.7 μM), *ndc-1*guide 2 (11.7 μM), purified Cas9 protein (qb3 Berkeley, 14.7 μM), *dpy-10* guide (3.7 μM) and a *dpy-10* repair template (29 ng/mL). [37, 38]

The RNA-protein mix was then injected into the gonads of N2 adult worms, which were then allowed to recover for three days. After this recovery, F1 progeny were screened for a roller phenotype and 10-20 F1s were singled out to individual plates. These roller worms were allowed to produce progeny, which were then genotyped by PCR to detect the presence of the *npp- 22/ndc1* deletion allele. The deletion strain was outcrossed four times to N2 (ancestral strain) worms before use and characterization.

#### CRISPR-Cas9 with SEC repair template for endogenous tagging

To endogenously tag *ndc-1* with mNeonGreen and mRuby, a self-excising cassette (SEC) repair template approach was utilized. 950Bp homology arms from the five- and three prime-end of *ndc1*’s stop codon were cloned into an SEC vector. This plasmid was co-injected with a Cas9 +ndc1 guide plasmid into the gonads of adult N2 worms. Worms were rescued to individual plates and allowed to recover for three days at 20°C. Plates were screened for roller worms and positive plates were treated with approximately 200-300 μl of hygromycin B (20 mg/mL) and were allowed to recover for four-five days. Plates that had surviving rolling worms were screened for mNEONgreen or mRuby signal by microscopy. Finally, worms were outcrossed four times to N2 worms, before crossing to other fluorescent markers.

## METHOD DETAILS

### RNAi

Primers were designed to amplify a 200-1000bp region within a gene of interest, see key resource table for list of primers used. Primers were designed to amplify from within a single exon whenever possible. N2 gDNA or cDNA was used as a template for PCR, which in turn was purified and used in T3/T7 reverse transcription reactions (MEGAscript, Life Technologies). The synthesized RNAs were purified using phenol-chloroform and resuspended in 1X soaking buffer (32.7 mM Na2HPO4, 16.5 mM KH2PO4, 6.3 mM NaCl, 14.2 mM NH4Cl). RNA reactions were annealed at 68°C for 10 minutes followed by 37°C for 30 minutes. dsRNAs were brought to a final concentration of ∼2000 ng/μl whenever possible and 2-μl aliquots of the dsRNA were stored until use at –80°C. For each experiment, prior to injection, a fresh aliquot was diluted to ∼1000ng/μl and centrifuged at 13,000 rpm for 30 minutes at 4°C. 0.35μl of the diluted dsRNA was loaded into the back of hand pulled capillary needles and injected into the gut of L4 worms. Worms were rescued to plates seeded with OP-50 and allowed to recover for ∼24 hours prior to imaging or lethality analysis.

### Lethality quantifications

L4 worms were injected with indicated dsRNA and allowed to recover for 24 hours at 20°C. 24 hours post-injection worms were then singled out and allowed to self-fertilize for an additional 24 hours. Worms were transferred to another plate for a final 24 hours, then disposed of. Plates corresponding to 24-48 hours and 48-72 hours post-injection were then scored for hatched larvae and unhatched embryos. Prior to counting, embryos were given 24 hours to hatch. The total number of embryos and larvae were combined for each time window to calculate embryonic lethality and brood size. A similar approach was used for noninjected control worms and worms containing deletion alleles.

### Immunoblots

#### Generation of whole worm lysate

For a given condition, a microcentrifuge tube was filled with 30 μl of M9 Buffer and the fill line was marked with a black marker. For each condition 35 adult worms were then placed in the microcentrifuge tube and washed three times with M9+0.1% Triton. After the final wash tubes were brought up to a final volume of 30 μl. Then, 10 μl of 4x sample buffer was added and the tubes were mixed. The samples were then sonicated at 70°C for 15 minutes, followed by incubation for five minutes at 95°C. Samples were re-sonicated at 70°C for an additional 15 minutes. Worm lysates were stored at -20°C until they were run on an SDS-PAGE protein gel.

#### Protein gel electrophoresis and antibody probing

For all protein gels, homemade 8%-10% SDS-PAGE were used. Worm lysates were re-boiled at 95°C for five minutes and then 20 μl (∼17.5 worms) were loaded into each lane. The protein gel was then run at 80V for 15 minutes to fully collapse samples. The protein gel was then run at 120V for approximately 90 minutes, or until the sample buffer reached the bottom of the gel. Protein samples were then transferred overnight (16 hours at 4°C) to an nitrocellulose membrane at 100mA. Membranes were blocked in fresh PBST for one hour at room temperature; membranes were then cut based on size and incubated overnight with the following primary antibodies: 1 μg/mL mouse α-alpha-tubulin (EMD Millipore), 1 μg/mL mouse mAb414 (Biolegend), rabbit α-NPP-5/Nup107 (1:300), rabbit α-NPP-10N/Nup98 (1:500), rabbit α-NPP-10C/Nup96 (1:500), and rabbit α-NPP-19/Nup53 (1:1000).

Membranes were then rinsed three times quickly with PBST followed by four five-minute washes. Membranes were then incubated with appropriate secondary antibodies for one hour at room temperature. Secondary antibodies were diluted 1:5000 for horseradish peroxidase (HRP)- conjugated goat-anti-rabbit and 1:7000 for HRP-conjugated goat-anti-mouse (Thermo Fischer Scientific). Membranes were rinsed and washed as described above. Prior to image acquisition membranes were incubated with Clarity Max Western ECL Substrate (BIO-RAD) for four minutes and then excess reagent was removed.

### Immunofluorescence

#### Slide preparation

Microscope slides (Fisher Scientific Premium Microscope Slides Superfrost) were coated with 0.1% polylysine and dried on a low temperature heat-block. Slides were then baked at 95 °C for 30 minutes. Slides were used the same day that they were baked.

#### Fixation and primary antibody incubation

15-20 adult worms were picked into a 4 μl drop of ddH_2_O and covered with a standard 18×18mm coverslip. Embryos were pushed out of the adult worms by pressing down on the corners of the coverslip with a pipet tip. To crack the eggshell and permeabilize the embryos, slides were placed in liquid nitrogen for ∼five-minutes. Coverslips were quickly removed by using a razor blade to pop off the coverslip. Slides were then fixed in pre-chilled 100% methanol at -20 °C for 20 minutes. Following fixation slides were washed two times in 1X PBS+ 0.2% Tween 20 (PBST) at room temperature for 10 minutes each. After the second wash, samples were blocked with 50 µL of 1 % BSA in PBST per slide in a humid chamber for one hour at 20°C. Slides were then incubated overnight at 4°C with primary antibodies diluted in PBST (45 μl per slide; rabbit α-LMN1, 1 μg/mL, mouse mAb414 2.5 μg/mL, rabbit α-NPP-5/Nup107 (1:300), rabbit α-NPP-19/Nup53 (1:300), and rabbit α-MEL-28/Elys (1:500).

#### Secondary antibody incubation and DAPI +Hoechst staining

After overnight primary antibody incubation, slides were washed two times in (50 μl per slide) PBST at room temperature for 10 minutes each. Following the second wash, slides were incubated at 20°C for one hour in the dark with secondary antibodies diluted in PBST (45 μl per slide, anti-rabbit Cy3/Rhodamine, 1:200; anti-mouse FITC, 1:200; [Jackson Immunoresearch]). Slides were again washed two times in (50μl per slide) PBST at room temperature for 10 minutes each in the dark. Samples were stained with 1 µg/mL Hoechst (diluted from a 1 mg/mL stock in H_2_O). Slides were washed two final times in (50μl per slide) PBST at room temperature for 10 minutes each in the dark. Finally, mounting media (Molecular Probes ProLong Gold Antifade Reagent with DAPI) was added to each sample. Coverslips were gently placed onto the slides and adhered with clear nail-polish. Slides were allowed to dry at 20°C and then stored at –20°C until they were imaged.

### Image acquisition

#### Live microscopy

First, 2% agarose imaging pads were made using molten agarose (95°C) on a glass slide. Adult hermaphrodites were then dissected on glass slides and the embryos were transferred to two percent agarose imaging pads using a mouth pipette at (∼20°C). Embryos were organized using an eyelash tool to group similar stage embryos. Images were acquired every 20 seconds. Five z-slices were taken for each time point with 2-μm steps. Images were acquired on an inverted Nikon Ti microscope equipped with a confocal scanner unit (CSU-XI, Yokogawa). Two solid state lasers (100-mW 488-nm and 50-mW 561-nm) were used in conjunction with a 60x objective lens (Å∼ 1.4 NA pan Apo). Images were recorded with a high-resolution ORCA R-3 Digital CCD Camera (Hamamatsu).

#### Fluorescece recovery after photobleaching (FRAP)

First, 2% agarose imaging pads were made using molten agarose (95°C) on a glass slide. Adult hermaphrodites were then dissected on glass slides and two-cell stage embryos were transferred to two percent agarose imaging pads using a mouth pipette at (∼20°C). Embryos were organized using an eyelash tool to group similar stage embryos. A stimulation ROI was then drawn on a region of the nuclear envelope of an AB nucleus (two cell stage embryo). Three images were taken prior to stimulation/bleaching of the ROI by a 100 nW, 405 nm laser. Images were then acquired for the remainder of the cell cycle, until nuclear envelope breakdown. Images were taken every 10s with 1 z-slice taken per time point.

#### Fixed microscopy

Immunofluorescent images were acquired on an inverted Nikon Ti Eclipse microscope. This microscope was equipped with solid state 100 mW 405, 488, 514, 594, 561, 594, and 640 nm lasers, a Yokogawa CSU-W1 confocal scanner unit, a 60x 1.4 NA plan Apo objective lens, and a prime BSI sCMOS camera.

### qPCR

For each strain (10-15 medium plates) unsynchronized populations of adult worms were washed with 1x M9 buffer into 15-ml conical tubes. Worms were washed with five mL of M9, three times. After the final wash, the buffer was removed from the worm pellet and the samples were frozen at –80°C. Next an approximate 100 µl worm pellet was ground using a motorized micro pestle. 1 mL of Trizol was then added to the ground worm pellet and the mixture was vortexed for 15 minutes at room temperature. Following vortexing, 200µl of chloroform was added to the sample. The trizol-chloroform mixture was then briefly vortexed (15s) followed by an incubation at room temperature for five minutes. Next the solution was spun for 15 minutes at 4°C (12,000 rpm). To extract the total RNA, the upper aqueous layer was removed (approximately 500µl) and transferred to a new microcentrifuge tube. 500 µl of isopropanol was added to the tube and it was inverted six times.

To precipitate the RNA, the solution was incubated at room temperature for 10 minutes followed by a 10-minute spin 4°C (12,000 rpm). The supernatant was discarded, and the pellet was washed in 1 mL of 75% ethanol by inverting once and vortexing for 10s. To pellet the RNA, the solution was spun for 5 minutes at 4°C (7500 rpm), the supernatant was removed, and the pellet was air-dried at room temperature for 5 minutes. The pellet was resuspended in 87 µl of RNase-free water and incubated at 37°C for 15 minutes. After incubation, the isolated RNA was mixed with 10 µl NEB Buffer 4, 1 µl of 50 µM CaCl2, and 2 µl Ambion Turbo DNase and incubated at room temperature for 15 minutes. The RNA/DNase reaction was cleaned with a phenol-chloroform extraction and eluted in 20 µl of RNase free water. 500ng of total RNA was reverse transcribed using Invitrogen Super-Script II Reverse Transcriptase kit. The cDNA was stored at –20°C.

For qPCR reactions, cDNA (∼ 1200 ng/μl) was diluted fivefold and pre-pared for amplification using a BioRad SYBR Green Supermix Kit with the primers for nucleoporin genes listed in key resource table as well as *hxk-2* primers from [35]. For each biological replicate (two wildtype and two *ndc1Δ*), four technical replicates were performed. Reactions were loaded into a 384-well plate for amplification. The reactions were amplified, and *Ct* values measured using a BioRad CFX 384 qPCR machine. An annealing temperature of 55°C was used for all genes. After amplification, *Ct* values for each nucleoporin gene were normalized to *hxk-2*, and the fold change relative to the control was calculated using the 2–(ΔΔ*Ct*) method (BioRad).

### Transmission Electron Microscopy

#### Sample preparation

Wild-type N2 and SBW83 *C. elegans* hermaphrodites were dissected in M9 buffer and single embryos early in mitosis were selected and transferred to cellulose capillary tubes (Leica Microsystems, Vienna, Austria) with an inner diameter of 200 μm. The embryos were observed with a stereomicroscope until cleavage furrow ingression in late anaphase and then immediately cryo-immobilized using a LEICA ICE high-pressure freezer (Leica Microsystems, Vienna, Austria). Freeze substitution was performed over 3 d at -90 °C in anhydrous acetone containing 1 % OsO_4_ and 0.1 % uranyl acetate using an automatic freeze substitution machine (EM AFS, Leica Microsystems, Vienna, Austria). Epon/Araldite infiltrated samples were flat embedded in a thin layer of resin, polymerised for 2 d at 60 °C, and selected by light microscopy for re-mounting on dummy blocks. Serial semi-thick sections (200 nm) were cut using an Leica Ultracut S Microtome (Leica Microsystems, Vienna, Austria). Sections were collected on Pioloform-coated copper slot grids and poststained with 2 % uranyl acetate in 70 % methanol followed by Reynold’s lead citrate.

#### Data acquisition by electron tomography

Colloidal gold particles (15 nm; Sigma-Aldrich) were attached to both sides of semi-thick sections collected on copper slot grids to serve as fiducial markers for subsequent image alignment. For dual-axis electron tomography, series of tilted views were recorded using an F20 electron microscopy (Thermo-Fisher, formerly FEI) operating at 200kV at magnifications ranging from [5000x-6500x], and recorded on a Gatan US4000 (4000px X 4000px) CCD or a Teitz TVIPS XF416 camera. Images were captured every 1.0° over a ±60° range.

#### 3D reconstruction and automatic segmentation of microtubules

We used the IMOD software package (http://bio3d.colourado.edu/imod), which contains all of the programs needed for calculating electron tomograms. For image processing the tilted views were aligned using the positions of the colloidal gold particles as fiducial markers. Tomograms were computed for each tilt axis using the R-weighted back-projection algorithm.

#### Nuclear envelope and NPC segmentation and measurement

The IMOD software package was used to segment the nuclear envelope and nuclear pore complexes in *cntrl* and *ndc1Δ* tomograms. Regions of continuous nuclear membranes were traced in the “non-core” region of the reforming nuclear envelope. Three criteria were used to distinguish nascent NPCs from simple NE holes, 1) the gaps between the two membrane edges were less than 100nm, 2) the two membrane edges tapered to a point suggesting there was fusion of the inner and outer NE, and 3) there was stretches of continuous membranes above and below the gap.

## QUANTIFICATION AND STATISTICAL ANALYSIS

### Image analysis

#### Nuclear import analysis

To determine the fluorescence intensity of NLS-LacI∷GFP inside the nucleus of one- and two-cell stage embryos, the chromatin was traced with either the freehand or circle tool in FIJI . Camera background was determined by drawing a 50×50 pixel box in vacant areas of the video. Average cytoplasmic values were determined by drawing a 20×20 pixel box inside the embryo. The nuclear to cytoplasmic ratio (N:C) was determined by subtracting the average camera background from each value and then the nuclear value was divided by the cytoplasmic value. To account for differences in nuclear size, this ratio was then multiplied by the nuclear area. This process was repeated for each time point and condition.

#### Line scan analysis of two-cell stage embryos

A three-pixel wide by 10-micron long line was drawn and centered on the nucleus to determine the fluorescence intensity. The line was then redrawn perpendicular to the first and a second measurement was taken. The values from these two lines were then averaged to give the fluorescence intensity of the nuclear envelope. The average value of the first and last 2 points of each line were used to subtract background from the rest of the line. These values were then plotted against the relative position along the line.

#### Line scan analysis of immunofluorescent images

A three-pixel wide by five-micron long line was drawn and centered on the nuclear envelope. The line was then redrawn perpendicular to the first and a second measurement was taken. The values from these two lines were then averaged to give the fluorescence intensity of the nuclear envelope. The same line scan was relocated to a clear area of the video to get the average intensity for camera background. Finally, to account for wide differences in fluorescence intensity, the data was normalized by dividing each value by the maximum fluorescence intensity. The final fluorescent intensities were then plotted against the relative position along the line. Additionally, the normalized value at the nuclear envelope was divided by the first five values on the line to determine the NE:cytoplasmic ratio (shown in Figure xx) or by the last five values to determine the NE:nucleoplasmic ratio (shown in Figure XX).

#### Quantification of Nup160:GFP Puncta and colocalization with Ndc1:mRuby

To determine the number of NPP∷GFP puncta/aggregates in the cytoplasm of one-cell stage embryos, maximum projections were generated using FIJI. The puncta were then tracked using the TrackMate plugin in FIJI [39]. This process was repeated to determine the number of puncta in *ndc1*, *npp-7* RNAi depleted embryos. To determine the colocalization between the Nup160:GFP puncta and Ndc1:mRuby puncta, three-pixel wide line scans were drawn in each channel and plots were overlayed

#### FRAP analysis

To evaluate the amount of recovery for Nup160:GFP and Ndc1:mNG, a rectangular ROI was used to measure the average fluorescence intensity of the nuclear envelope before and after photobleaching. These values were normalized to the pre-bleach frame and graphed relative to bleaching. To calculate the mobile/immobile fraction as well as the T1/2, the data was fit to the exponential equation f(t)=A*(1-exp(-tau*Time). To obtain the best fit, the parameters were fitted to the data using nonlinear least squares in R-studio.

### Statistical Analysis

All data points are reported in graphs and error bar types are noted in figure legends. Statistical analysis was performed on datasets with multiple samples and independent biological repeats. The type of test used, sample sizes, and P values are reported in figure legends or in text (P < 0.05 defined as significant). Statistical tests were performed using GraphPad Prism.

## Notes

### Competing Interest Statement

The authors have declared no competing interest.

